# *In vivo* phenotyping of Parkinson-specific stem cells reveals increased a-Synuclein levels but no spreading

**DOI:** 10.1101/140178

**Authors:** Kathrin Hemmer, Lisa M. Smits, Silvia Bolognin, Jens C. Schwamborn

## Abstract

Parkinson′s disease is a progressive age-associated neurological disorder. One of the major neuropathological hallmarks of Parkinson’s disease is the appearance of protein aggregates, mainly consisting of the protein alpha-Synuclein. These aggregates have been described both in genetic as well as idiopathic forms of the disease. Currently, Parkinson’s disease patient-specific induced pluripotent stem cells (iPSCs) are mainly used for *in vitro* disease modeling or for experimental cell replacement approaches. Here, we demonstrate that these cells can be used for *in vivo* disease modeling. We show that Parkinson’s disease patient-specific, iPSC-derived neurons carrying the LRRK2-G2019S mutation show an upregulation of alpha-Synuclein after transplantation in the mouse brain. However, further investigations indicate that the increased human alpha-Synuclein levels fail to induce spreading or aggregation in the mouse brain. We therefore conclude that grafting of these cells into the mouse brain is suitable for cell autonomous *in vivo* disease modeling but has strong limitations beyond that. Furthermore, our results support the hypothesis that there might be a species barrier between human to mouse concerning alpha-Synuclein spreading.

## Introduction

Parkinson′s disease is a progressive neurological disorder. It is the second most common neurodegenerative disease after Alzheimer’s disease. Parkinson’s disease is characterized by motor symptoms including tremor, rigidity, bradykinesia, and postural instability but also by non-motor symptoms as fatigue, depression, sleep disturbance, and dementia (Postuma et al., 2012). Although some symptomatic treatments exist, no preventive or disease-modifying therapies are currently available. The major hallmark of Parkinson’s disease is the degeneration of dopaminergic neurons in the substantia nigra of the midbrain. It is estimated that about 30 % of the familial and 3-5 % of the sporadic cases are caused by monogenetic mutations (Klein and Westenberger, 2012). The affected genes including *SNCA, PINK1, Parkin, LRRK2, DJ1, ATP13A2* and *VPS35* are involved in numerous, very different biochemical processes, emphasizing a complex etiopathogenesis. Among others, cellular processes like oxidative stress, dysfunctional protein degradation and clearance, calcium dysregulation, mitochondrial dysfunction as well as protein aggregation (prionopathy) have been implicated in the etiopathogenesis of Parkinson’s disease. Although the majority of the Parkinson’s disease cases is of idiopathic origin, the clinical symptoms and the histopathology do not differ from the genetic Parkinson’s disease cases (Klein and Westenberger, 2012). Therefore, to study genetic cases might serve as a tool to get more insights into the complex etiopathogenesis of the pathology. A major neuropathological hallmark of Parkinson’s disease is protein aggregates that are detectable in many genetic as well as idiopathic cases. These aggregates are called Lewy bodies (LB) and Lewy neurites and consist mainly of the protein alpha-Synuclein (gene: *SNCA*). Several mutations in *SNCA* as well as increased *SNCA* gene doses have been previously described to cause Parkinson’s disease (reviewed in (Kalinderi et al., 2016)). Interestingly, neuropathological studies suggest that Lewy pathology ascends from peripheral autonomic ganglia to brainstem nuclei and subsequently toward the neocortex over time (Braak and Del Tredici, 2009). The associated prionopathy hypothesis posits that conformationally altered alpha-Synuclein is transmitted between neurons and initiates protein aggregation in susceptible neurons (Braak and Del Tredici, 2009). This hypothesis is supported by the observation that alpha-Synuclein fibrils induce LB pathology in primary neuronal cultures as well as in wild-type mice (Luk et al., 2012a; Volpicelli-Daley et al., 2011). However, whether increased levels of alpha-Synuclein in a subset of cells would be sufficient to induce such a prion-like spreading remains unclear so far.

The reprogramming of human somatic cells into induced pluripotent stem cells (iPSCs) was a breakthrough for *in vitro* disease modeling (Takahashi and Yamanaka, 2006; Yu et al., 2007). iPSCs resemble embryonic stem cells in all their characteristics. Numerous studies used Parkinson’s disease patient-specific iPSCs and thereof-derived neurons to gain insights into the mechanisms underlying the onset and progression of Parkinson’s disease (Hillje and Schwamborn, 2016). Among other findings, it was possible to demonstrate that even mutations in Parkinson’s disease associated genes different from *SNCA*, can cause an upregulation of the alpha-Synuclein protein levels (Sanchez-Danes et al., 2012). However, the usage of human iPSC-derived cellular models under physiological conditions, e.g. via grafting in mice, still remains unexplored.

In this study, we used an isogenic pair of Parkinson’s disease patient-specific iPSCs with the G2019S mutation in the gene *LRRK2* and transplanted thereof-derived neuroepithelial stem cells (NESCs) into the striatum of mice. We demonstrated that *in vivo* differentiated neurons showed an upregulation of alpha-Synuclein under physiological conditions. However, we were unable to detect any spreading of alpha-Synuclein in the mouse brain. Our results suggest that human Parkinson’s disease patient-derived iPSC models were able to recapitulate key characteristics of the disease *in vivo*. Furthermore, they support the hypothesis that murine alpha-Synuclein might actually inhibit seeding and propagation of human alpha-Synuclein.

## Material and Methods

### Stem cell culture

Human induced pluripotent stem cells (iPSCs) were derived from an 81 old, female Parkinson’s disease patient carrying the LRRK2-G2019S mutation (Reinhardt et al., 2013b). The iPSC line was gene corrected by using Zinc Finger Nucleases (Reinhardt et al., 2013b).

From these iPSCs, neuroepithelial stem cell (NESC) lines were generated and cultured as described elsewhere (Reinhardt et al., 2013a). In brief, cells were cultured on Matrigel-coated plates in N2B27 medium (DMEM-F12 (Invitrogen)/Neurobasal (Invitrogen) (50:50), 1:200 N2 supplement (Invitrogen,), 1:100 B27 supplement w/o Vitamin A (Invitrogen) 1:100 penicillin/streptomycin (Invitrogen), 1:100 L-Glutamine (Invitrogen) freshly supplemented with 0.5 μM Purmorphamine (Enzo Life Science), 3 μM CHIR (Axon Medchem), and 150 μM ascorbic acid (Sigma). Cells were split 1:10-1:15 once per week using Accutase. Differentiation was initiated by changing medium two days after splitting to N2B27 medium containing 1 μM PMA, 200 μM AA, 10 ng/mL BDNF (Peprotech), 10 ng/mL GDNF (Peprotech), 1 ng/mL TGF-b3 (Peprotech), and 500 μM dbcAMP (Sigma).

### Transplantation

All animal experimentation was approved by the appropriate Luxembourg governmental agencies (Ministry of Health and Ministry of Agriculture). For surgeries 11-12 week old NOD.Cg-Prkdcscid Il2rgtm1Wjl/SzJ mice were deeply anesthetized by isoflurane (4 % v/v for induction and 2 % v/v for maintenance) and bupivacain (5 mg/kg s.c.) was given as supplementary local analgesia before surgery. Additionally, buprenorphine treatment (0.1 mg/kg; s.c.;) was used as analgesia before and one day after surgery. Two weeks before transplantation, 10 μg 6-OHDA (6-Hydroxydopamine hydrobromide, Sigma Aldrich) was stereotactically injected into the striatum using following coordinates in relation to bregma: anteroposterior: 0.5 mm, mediolateral: 2.0 mm, dorsoventral: 3.5 mm (below skull). Transplantation of NESCs was performed as described before (Hemmer et al., 2014; Reinhardt et al., 2013a). In brief, NESCs (Passage 10-13) were differentiated towards dopaminergic neurons for 6 days. For transplantation, cells were dissociated to single cells with Accutase and resuspended in N2B27 medium to a concentration of 5 × 10^4^ cells per microliter. Three microliters of the cell suspension was injected into the 6-OHDA-lesioned striatum using a Hamilton 7005KH 5-μl syringe (LRRK2-G2019S n=6, LRRK2-WT n=3).

### Perfusion, sectioning, and immunohistochemical analysis

Perfusion, sectioning, and immunohistochemical analysis were performed 11 weeks post-transplantation as described previously (Hemmer et al., 2014; Reinhardt et al., 2013a). The following primary antibodies were used: human Nuclei (mouse, 1:200, Millipore), human NCAM (mouse, 1:100, Santa Cruz), and alpha-Synuclein (rabbit, 1:600, Sigma). Alexa fluorophore-conjugated secondary antibodies (Invitrogen) and Hoechst 33342 (Invitrogen) were used to visualize primary antibodies and nuclei, respectively. Sections (40 μm) from the center of the graft were evaluated using a Zeiss LSM 710 confocal microscope. 3D surface structures of the z-stacks taken by the confocal microscope were created using IMARIS software (bitplane). For this, three different ROIs (graft, proximal to graft and distal to graft) were chosen and the values of the threshold (background substraction) were automatically set. The endogenous mouse stainings of the striatal myelinated fibers were not taken into consideration for the analysis.

### Data and statistical analysis

The volume, the mean intensity, and maximum intensity of every created surface were defined by Imaris software. The average or the sum of the different parameters was calculated for each animal. Ratios of the different ROIs were created as indicated and t-tests were performed using SigmaPlot. n specifies the number of mice and statistical significance is considered to be P < 0.05. Prism 6.01 (GraphPad) was used for data illustration and data are presented as mean + SEM.

## Results

### Parkinson’s disease-specific neurons, expressing LRRK2-G2019S, upregulated alpha-Synuclein *in vivo*

Previously, it has been shown *in vitro* that human neurons expressing the Parkinson’s disease-associated G2019S mutation in the *LRRK2* gene have elevated levels of alpha-Synuclein (Sanchez-Danes et al., 2012). As a first step, we were aiming at determining whether this is also the case *in vivo*. Accordingly, we transplanted 1,5 × 10^5^ human iPSC-derived NESCs that were pre-differentiated for the neuronal lineage (for details see Materials and Methods section), into the striatum of 11-12 week old NOD.SCID mice. In order to investigate the impact of the LRRK2-G2019S mutation, we used patient-derived cells expressing this mutation as well as a corresponding isogenic line where the mutation has been corrected. Human cells were distinguished from the surrounding mouse cells with human-specific antibodies against NCAM and human nuclei (Fig. 1A, B). 11 weeks after transplantation in the mouse striatum, we saw robust survival and neuronal differentiation for both lines as indicated with the anti-human-NCAM antibody (Fig. 1A). In order to detect the levels of alpha-Synuclein in the grafted cells, we stained with anti-alpha-Synuclein-specific antibodies (Fig. 1B). For quantification, we reconstructed the surfaces of the alpha-Synuclein signal in the grafts with IMARIS (Fig. 1C). The volume and the intensity of the alpha-Synuclein signal was quantified from the reconstruction (Fig. 1D, E). This quantification revealed that expression of LRRK-G2019S led to a negligible increase in the mean volume of the alpha-Synuclein signal (Fig. 1D). However, the mean and the maximum intensity of the alpha-Synuclein signal was significantly increased upon expression of the Parkinson’s disease-associated LRRK2-G2019S mutation (Fig. 1E). These results indicate that human neurons carrying the LRRK2-G2019S mutation express higher levels of alpha-Synuclein under physiological conditions.

**Figure 1.**
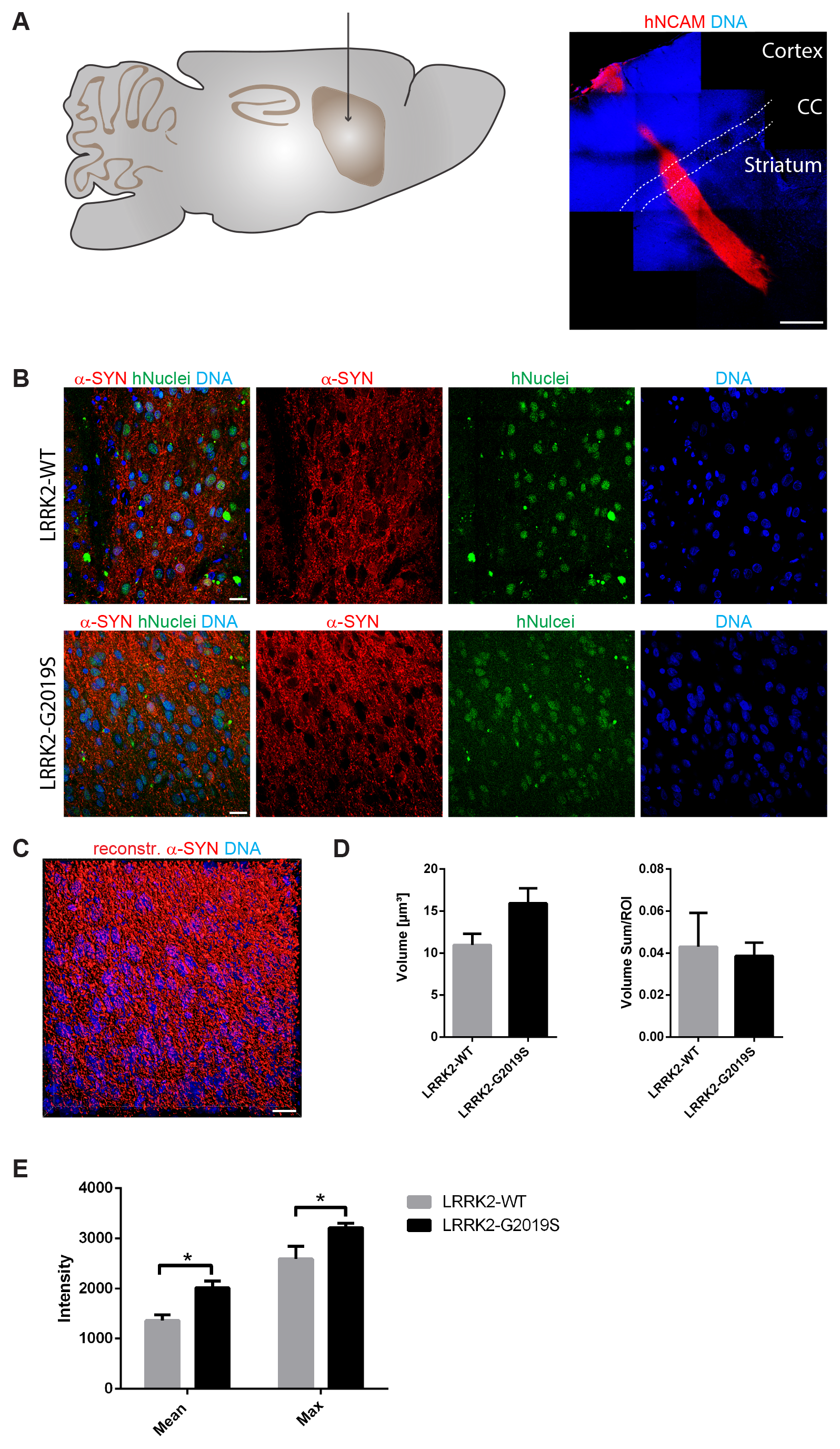
LRRK2-G2019S mutation led to a higher expression of alpha-Synuclein in grafted neurons. **A** Schematic overview of the adult mouse brain (left) representing the stereotactic target side of the striatum for the transplantation. 11 weeks after transplantation, the species-specific antibody hNCAM revealed a sound survival and neuronal differentiation of the graft (right). Dashed lines indicate the corpus callosum. **B** Immunohistological stainings indicated the expression of alpha-Synuclein in both cell lines after transplantation. **C** Representative 3D surface reconstruction of alpha-Synuclein of the graft shown in the lower panel of B. **D** Quantification of the volume and volume sum/region of interest of the reconstructed alpha-Synuclein surfaces. **E** Quantification of the mean and maximum intensity of the reconstructed alpha-Synuclein surfaces. Scale bars: 500 μm (A) and 20 μm (B and C). Error bars represent mean + SEM (LRRK2-WT n=3, LRRK2-G2019S n=6); α-SYN: alpha-Synuclein, CC: corpus callosum, max: maximum, ROI: region of interest

### Grafted neurons did not induce alpha-Synuclein spreading in the mouse brain

Parkinson’s disease is a progressive disorder characterized by the spreading of protein aggregates, mainly consisting of alpha-Synuclein. It has been hypothesized that this appearance of aggregates spreads in a prion-like fashion (Braak and Del Tredici, 2009). Aggregated alpha-Synuclein has the ability to induce further aggregation of soluble alpha-Synuclein. Hence, a seed of alpha-Synuclein aggregation would be sufficient to start the process.

In order to test whether our *in vivo* phenotyping approach would be able to recapitulate this spreading process, we analyzed the levels of endogenous mouse alpha-Synuclein in a region close (proximal) to the graft in comparison to a region far (distal) from the graft (Fig. 2A). Accordingly, a 3D surface reconstruction of the endogenous mouse alpha-Synuclein signal was performed with IMARIS (Fig. 2B and C). If alpha-Synuclein indeed would spread from the graft into the surrounding tissue, we expected to find significant differences in the volume (Fig. 2D) or intensity (Fig. 2E, F) of the alpha-Synuclein signal in a region proximal to the graft compared to a region distal to the graft when normalizing both regions to the graft. Particularly, we expected to see such differences in cases where cells with the LRRK2-G2019S mutation have been transplanted. Moreover, we compared the ratio of the two regions between the different genotypes. However, in none of the investigated parameters significant differences were detectable (Fig. 2D-F). We therefore conclude that in the here chosen paradigm Parkinson’s disease-associated alpha-Synuclein spreading is not detectable.

**Figure 2.**
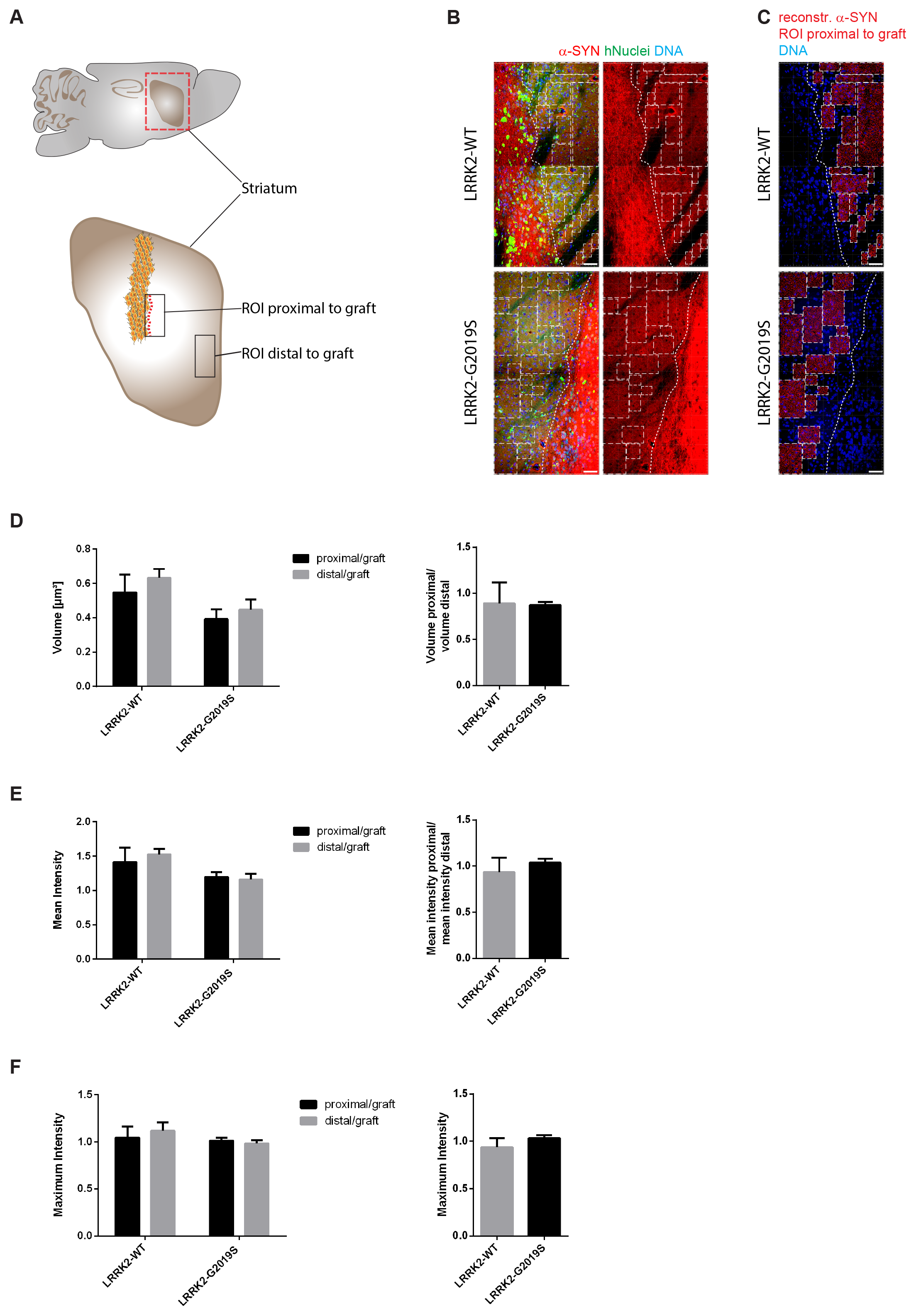
Alpha-Synuclein spreading from the graft was not detected in the surrounding tissue. **A** Schematic overview of the mouse adult brain indicating the ROIs that were chosen to analyse endogenous mouse alpha-Synuclein close (proximal) as well as far (distal) from the graft. **B** Representative images of the ROIs that were chosen proximal to the graft. Dashed lines define the edges of the graft. Squares mark the ROIs proximal to the graft that were used to create 3D surfaces shown in C. **C** 3D surface reconstruction of endogenous mouse alpha-Synuclein of the ROIs shown in B. **D**-**F** Quantification of the volume, the mean intensity and the maximum intensity of the reconstructed endogenous mouse alpha-Synuclein surfaces proximal to the graft and distal to the graft. Left: Proximal and distal ROIs of endogenous mouse alpha-Synuclein were normalized to the ROIs of the graft and eventually the different distances were compared to each other within the same cell line. Right: The ratio of the different regions of endogenous mouse alpha-Synuclein were compared between the different cell lines. Scale bars: 50 μm (B and C). Error bars represent mean + SEM (LRRK2-WT n=3, LRRK2-G2019S n=6); α-SYN: alpha-Synuclein, max: maximum, ROI: region of interest

## Discussion

The spreading of alpha-Synuclein protein aggregates, in a prion-like fashion, is believed to be the underlying mechanism for the progression of Parkinson’s disease pathology through the human brain (Braak and Del Tredici, 2009). Furthermore, somatic mutations occurring during embryogenesis can lead to genetic mosaicism in the brain. Typically, genotyping is conducted in mesoderm-derived lymphocytes. Therefore, mutations in ectoderm-derived neural cells will be missed (Proukakis et al., 2013). Consequently, it is conceivable that Parkinson’s disease patients that are classified as idiopathic, actually carry a Parkinson’s disease-associated mutation in a subset of neurons. In fact, somatic mutations have been previously associated to neurodegenerative disorders. It was suggested that differences in Parkinson’s disease phenotype in monozygotic twins with LRRK2 mutations are caused by additional somatic mutations (Schneider and Johnson, 2012; Van Broeckhoven, 2010). In the past, addressing the hypothesis of prion-like spreading of alpha-Synuclein and genetic mosaicism as potential source for aggregates of alpha-Synuclein was complicated because of the lack of appropriate models. In this study, we show that Parkinson’s disease patient-specific iPSC-derived neurons upregulated alpha-Synuclein after transplantation which is believed to be a first important step to induce protein aggregation and spreading. Therefore, we conclude that this approach can be used for *in vivo* disease modeling. However, we further demonstrate that this alpha-Synuclein failed to spread in the mouse brain.

At first glance, it seems surprising not to see alpha-Synuclein spreading, because previous studies *in vitro* (Luk et al., 2009; Volpicelli-Daley et al., 2011) as well as *in vivo* (Luk et al., 2012a; Luk et al., 2012b) have shown spreading. Furthermore, our finding that Parkinson’s disease-specific cells with the LRRK2-G2019S mutation showed higher levels of alpha-Synuclein, supports their usability for modeling Parkinsons disease *in vivo* and indicates that they in principle might be able to induce alpha-Synuclein aggregation and spreading. However, it is noticeable that all these studies did not use a cross-species approach. Either human alpha-Synuclein spreading was investigated in human cells or in mice engineered to express human alpha-Synuclein. On the other hand, mouse alpha-Synuclein was tested in mouse cells or mice *in vivo*. Interestingly, a recent study using a cross-species approach reported that the mouse-alpha-Synuclein protein significantly attenuated the formation of aggregates of human-alpha-Synuclein (Fares et al., 2016). In other words, the mouse-alpha-Synuclein inhibited the aggregation of the human form, which implies the existence of a species barrier. This finding is also supported by a report showing that mouse-alpha-Synuclein inhibited the fibrillization of human-alpha-Synuclein *in vitro* (Rochet et al., 2000). In this context, it is further interesting to note that the A53T mutation in human-alpha-Synuclein causes Parkinson’s disease while the mouse version of alpha-Synuclein naturally expresses a Threonine at position 53. However, based on the here obtained results we cannot rule out that the failure to see alpha-Synuclein spreading in the mouse brain could be due to initially too low levels of human alpha-Synuclein in the transplanted cells. Additionally, we cannot exclude that an even longer duration of the experiment would have led to mouse alpha-Synuclein upregulation, aggregation, and spreading. Finally, the depletion of dopaminergic neurons in the striatum with 6-OHDA, preceding the transplantation, might have had a negative impact on the potential spreading. However, overall these results clearly indicate that it is of critical importance to choose the appropriate model and tools to study alpha-Synuclein aggregation and spreading. In particular, new human-specific organoid models recapitulating essential features of the human midbrain (Jo et al., 2016; Monzel et al., 2017) might represent interesting alternatives for the future. These models will allow combining the advantages of tissue-like specimens (complex system, *in vivo*, cellular heterogeneity) with high relevance for the human situation, particularly when they are derived from human Parkinson’s disease patient-specific stem cells.

## Abbreviations

α-SYN: alpha-Synuclein,
CC: corpus callosum,
iPSCs: induced pluripotent stem cells,
LB: Lewy bodies,
max: maximum,
NESCs: neuroepithelial stem cells,
ROI: region of interest

## Acknowledgments

The authors would like to thank Inga Brüggemann and Thea van Wüllen for excellent technical assistance. We further acknowledge support through the pluripotent stem cell facility at the LCSB. The JCS lab is supported by the Fonds National de la Recherche (FNR) (CORE, C13/BM/5791363). This is an EU Joint Programme. – Neurodegenerative Disease Research (JPND) project (INTER/JPND/14/02; INTER/JPND/15/11092422). Further support comes from the SysMedPD project which has received funding from the European Union’s Horizon 2020 research and innovation program under grant agreement No 668738. KH received financial support from a private philanthropist as well as from the Fondation du Pélican de Mie et Pierre Hippert-Faber. LMS is supported by a fellowship from the FNR (AFR, Aides à la Formation-Recherche).

## References

Braak, H., Del Tredici, K., 2009. Neuroanatomy and pathology of sporadic Parkinson’s disease. Adv Anat Embryol Cell Biol. 201, 1–119.

Fares, M.B., et al., 2016. Induction of de novo alpha-synuclein fibrillization in a neuronal model for Parkinson’s disease. Proc Natl Acad Sci U S A. 113, e912–21.

Hemmer, K., et al., 2014. Induced neural stem cells achieve long-term survival and functional integration in the adult mouse brain. Stem Cell Reports. 3, 423–31.

Hillje, A.L., Schwamborn, J.C., 2016. Utilization of stem cells to model Parkinson’s disease – current state and future challenges. Future Neurology. 11, 171–186.

Jo, J., et al., 2016. Midbrain-like Organoids from Human Pluripotent Stem Cells Contain Functional Dopaminergic and Neuromelanin-Producing Neurons. Cell Stem Cell. 19, 248–57.

Kalinderi, K., Bostantjopoulou, S., Fidani, L., 2016. The genetic background of Parkinson’s disease: current progress and future prospects. Acta Neurol Scand. 134, 314–326.

Klein, C., Westenberger, A., 2012. Genetics of Parkinson’s disease. Cold Spring Harb Perspect Med. 2, a008888.

Luk, K.C., et al., 2009. Exogenous alpha-synuclein fibrils seed the formation of Lewy body-like intracellular inclusions in cultured cells. PNAS. 106.

Luk, K.C., et al., 2012a. Pathological alpha-synuclein transmission initiates Parkinson-like neurodegeneration in nontransgenic mice. Science. 338, 949–53.

Luk, K.C., et al., 2012b. Intracerebral inoculation of pathological alpha-synuclein initiates a rapidly progressive neurodegenerative alpha-synucleinopathy in mice. J Exp Med. 209, 975–86.

Monzel, A.S., et al., 2017. Derivation of human midbrain-specific organoids from neuroepithelial stem cells. Stem Cell Reports. 8.

Postuma, R.B., et al., 2012. How does parkinsonism start? Prodromal parkinsonism motor changes in idiopathic REM sleep behaviour disorder. Brain. 135, 1860–70.

Proukakis, C., Houlden, H., Schapira, A.H., 2013. Somatic alpha-synuclein mutations in Parkinson’s disease: Hypothesis and preliminary data. Movement Disorders. 28, 705–712.

Reinhardt, P., et al., 2013a. Derivation and expansion using only small molecules of human neural progenitors for neurodegenerative disease modeling. PLoS One. 8, e59252.

Reinhardt, P., et al., 2013b. Genetic correction of a LRRK2 mutation in human iPSCs links parkinsonian neurodegeneration to ERK-dependent changes in gene expression. Cell Stem Cell. 12, 354–67.

Rochet, J.C., Conway, K.A., Lansbury, P.T., Jr., 2000. Inhibition of fibrillization and accumulation of prefibrillar oligomers in mixtures of human and mouse alpha-synuclein. Biochemistry. 39, 10619–26.

Sanchez-Danes, A., et al., 2012. Disease-specific phenotypes in dopamine neurons from human iPS-based models of genetic and sporadic Parkinson’s disease. EMBO Mol Med. 4, 380–95.

Schneider, S.A., Johnson, M.R., 2012. Monozygotic twins with LRRK2 mutations: Genetically identical but phenotypically discordant. Movement Disorders. 27, 1203–1204.

Takahashi, K., Yamanaka, S., 2006. Induction of pluripotent stem cells from mouse embryonic and adult fibroblast cultures by defined factors. Cell. 126, 663–676.

Van Broeckhoven, C., 2010. The future of genetic research on neurodegeneration. Nat Med. 16, 1215–7.

Volpicelli-Daley, L.A., et al., 2011. Exogenous alpha-synuclein fibrils induce Lewy body pathology leading to synaptic dysfunction and neuron death. Neuron. 72, 57–71.

Yu, J., et al., 2007. Induced pluripotent stem cell lines derived from human somatic cells. Science (New York, N.Y.). 318, 1917–1920.

